# Incorporating Prior Genomic Dose-Response Data to Support the Benchmark Dose Estimation of Toxicogenomics

**DOI:** 10.1101/2022.07.20.500899

**Authors:** Chao Ji, Kan Shao

## Abstract

Chemical risk assessment is an important tool to evaluate the toxicity of chemicals in the environment, and high throughput toxicogenomics plays an increasingly important role in risk assessment. In toxicogenomics, dose-response analysis for each gene is a data-limited situation, and thus parameter and benchmark dose (BMD) estimations typically have large uncertainty. To solve this problem, an informative prior by synthesizing toxicological information is integrated into the Bayesian benchmark dose modeling system (BBMD), a leading web-based toxicogenomics analysis application. We analyzed 276,126 toxicogenomics dose-response datasets and obtained plausible estimation of informative priors for seven commonly used continuous dose-response models. The effects of informative priors are investigated at the individual probe and pathway levels. Simulation studies based on six “true” models generated from typical genomic dose-response shapes show a significant decrease in uncertainty and an increase in accuracy of BMD estimates for most scenarios with informative priors than the counterpart with uninformative priors. The case study on the pathway analysis indicates that informative priors slightly improve the correlation between the pathway-based point of departure and apical point of departure. Overall, our study provides a practical strategy to incorporate existing toxicogenomic information as priors to improve the quality of chemical risk assessment.

**Graphic abstract:** 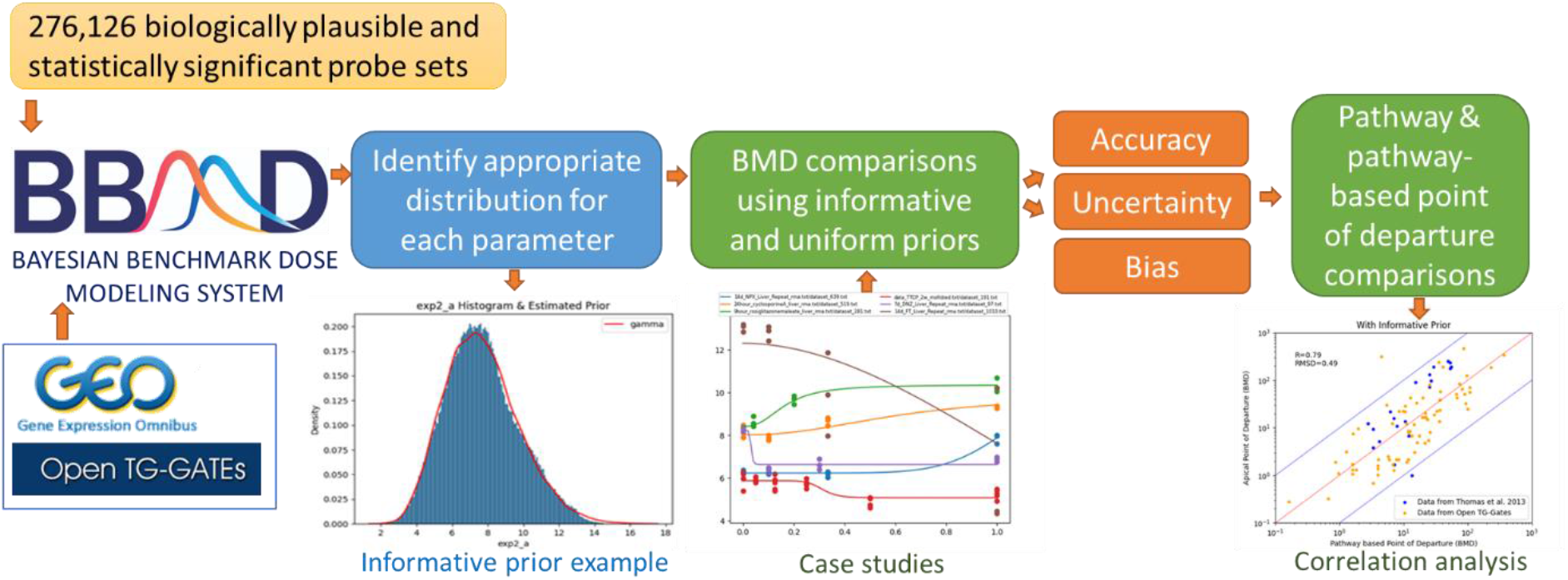

## 1. Introduction

Chemical risk assessment is widely applied in industries and regulatory agencies as an important tool to evaluate chemical toxicity in support of chemical registration, safety registration, and exposure limitation development. The benchmark dose (BMD) methodology is standard practice for the toxicity evaluation of chemicals ^4-5, 8^. As the histological animal toxicology experiments are time and resources consuming, there is a disparity between the capability of assessing chemicals by regulatory agencies and the annual increase of new chemicals^18^. Therefore, National Toxicology Program (NTP) proposed the initiative to apply toxicogenomics to predict toxicology ^13^. Studies have shown that toxicogenomics, which incorporates the genomic dose-response data to derive the in vivo pathway-based transcriptional biological effect points of departure (BEPOD), have the potential to facilitate the practice of chemical risk assessment ^1, 11, 17-18^. Recent developments in the toxicogenomics dose-response modeling systems ^6, 10, 14-15^ have made the practice of toxicogenomics feasible.

Toxicogenomics typically measures the responses of tens of thousands of probes in a microarray chip across control and several increasing doses. For each probe, the measurements are limited. Typically, 4-6 dose levels with three repeats are measured. For example, one of the most comprehensive datasets, Open TG-Gates^9^, has 4 dose levels and 3 repeated measurements for each tested probe. The commonly used continuous dose-response models have 2-4 parameters. Thus, estimated parameters and BMDs are typically associated with large variance (i.e., large uncertainty in estimation). The incorporation of appropriate informative priors could reduce the uncertainty^12^ in parameter and BMD estimates. One way to identify the appropriate informative priors is through the analysis of historical toxicological information^16^. However, currently, there are few studies to investigate ways to incorporate the historical data as informative priors for toxicogenomics and to reveal the effect of empirical data-derived informative priors on the precision and accuracy of toxicogenomics estimates. One reason is that the majority of toxicogenomics modeling system uses the maximum likelihood estimates, which could only assimilate the observations through likelihood function without the consideration of priors. On the contrary, the probabilistic Bayesian framework provides a useful tool to incorporate historical data as priors to enhance the robustness of dose-response modeling. Together, prior and observations as the likelihood function construct a posterior distribution for parameter estimation. On the other hand, informative priors might not be preferred because informative priors can lead to systematic biases and enlarged uncertainty^12^. But the effects of informative priors on the toxicology estimates using toxicogenomics are unknown. Hence, this paper aims to fill this knowledge gap and to understand the appropriate informative priors of toxicogenomics and its effects on BMD and BEPOD estimates on the ground of our previously developed Bayesian BMD (BBMD) estimation system for genomic data^10^.

The objective of this paper is to (1) determine appropriate empirical data-derived informative priors for common dose-response models, (2) investigate the impact of the priors on the precision and accuracy of each probe, and (3) examine the effects of informative priors on the correlation between BEPODs and their corresponding PODs.

## 2. Methods

In total, 276,126 biologically plausible and statistically significant probe sets that passed the one-way ANOVA preprocessing step with the default of fold change 2 and P-value 0.05 are kept deriving the informative priors. These statistically significant probe sets were collected from 1197 in vivo genomic dose-response datasets for 133-156 molecules of chemicals at different time scales (e.g., 3h, 6h, 9h, 24h, 4-5d, 1w, 2w, 4w, 13w) published in Open TG-Gates^9^ and NCBI’s Gene Expression Omnibus^3^ databases. The Open TG-Gates database was chosen because it is one of the most comprehensive toxicogenomic databases^2^ and it has both the in-vivo and histological data of the same chemical for comparison. These data are input to BBMD with vague uninformative priors which are all uniform distributions described in equations [S1]-[S7] in the Supplemental Material, and the lower and upper bounds of these uniform distributions are determined based on biological considerations and preliminary testing. The uniform distributions are used to cover all possible values as no parameter values are preferred when there is no information on the potential parameter value. In BBMD, for the dose-response data in each experiment, doses are normalized to 0-1 by dividing all dose levels by the highest dose. The normalized doses with associated responses are used for model fitting and these fitted parameters are used to derive informative priors of the commonly used continuous dose-response models. In these models, the power parameter *g* in the Hill, Power, Exponential 3, and Exponential 5 models in BBMD is recommended to be restricted to ≥1 in the BMD Technical Guidance^7^ to avoid a very steep slope in the low dose region resulting extremely low BMD estimates. However, the purpose of the prior derivation step is to reveal the density distribution of parameters among genes without restriction (i.e., *g* ≥ 1) that might change the shape of the distribution, so the power parameter in these models is set to be ≥0. The median value of the posterior sample is kept as the estimate of parameter for each dose-response dataset of probe sets, and is collectively being used to form histograms and derive informative priors. After the informative priors were derived, the power parameter was set to ≥1 to be consistent with dose-response modeling practice with genomic data^13^ for BMD and BEPOD calculations and comparisons. The pipeline of the analysis, shown in Figure 1, basically includes five steps: (1) data collection, (2) data assimilation with default uniform distribution, (3) informative prior derivation and incorporation to the BBMD system, (4) case study with six ‘true’ dose-response models for BMD estimation comparison, and (5) case study with published genomic data for BEPODs comparisons. Detailed steps on the prior analysis are described in Section 3 and the corresponding results follow thereafter.

**Figure 1.**
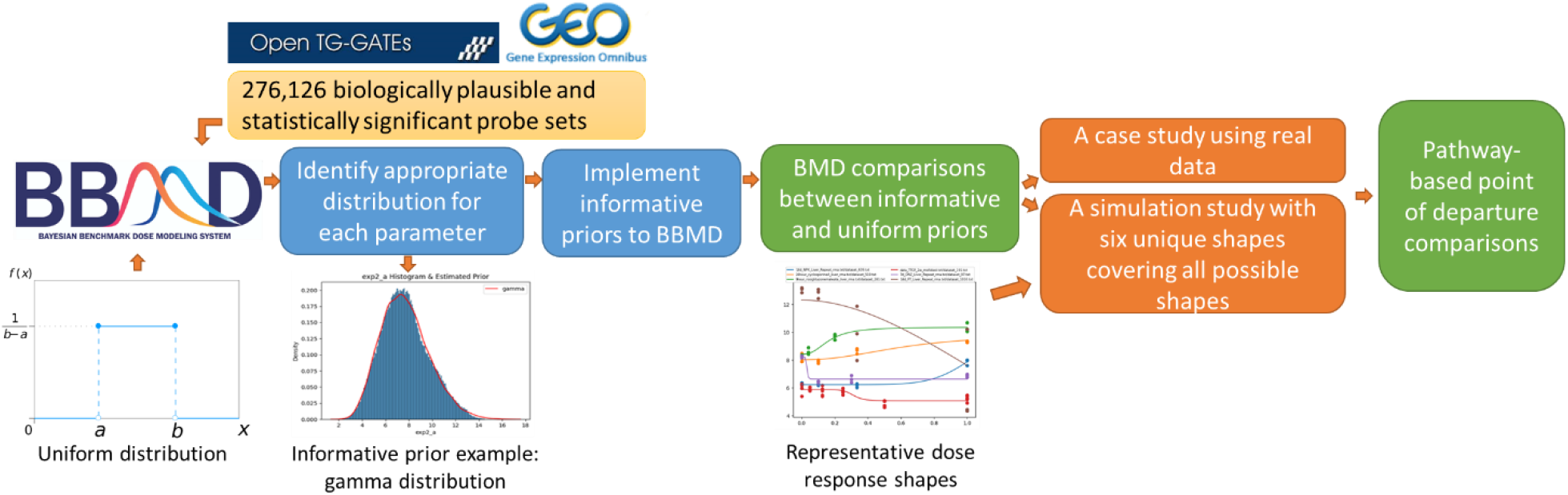
Pipeline of deriving informative priors

## 3. Analysis and results

### 3.1 Prior distributions estimate

#### 3.1.1 Prior analysis method

The dose levels of 276,126 probes are normalized and input in BBMD getting parameters estimates. The median values of the 21 parameters in seven models for 276,126 probes are plotted in histograms in Figures S1-S4. For distributions with a single peak, the ‘fitter’ python package is applied to identify the proper distributions among commonly used distributions including ‘Cauchy’, ‘chi-square’, ‘exponential’, ‘gamma’, ‘lognormal’, ‘normal’, and ‘beta’. For distributions with double peaks, a Gaussian mixture model is used. The candidate distribution is selected based on the following criteria (1) the parameter values should be consistent with the dose-response curve reflected by toxicological data (e.g., a slope parameter can be positive or negative to reflect that the slope can be increasing or decreasing) and (2) its shape should appropriately describe the shape of histograms. After the informative prior type is determined, the model parameters of the informative priors are estimated in the framework of a Bayesian hierarchical model. Here we take the linear model *y* = *a*_*j*_*x* + *b*_*j*_ (*y* is the response at dosage *x*) as an example to illustrate the Bayesian hierarchical framework in eq (1)-(2). The parameters are estimated using PyStan.

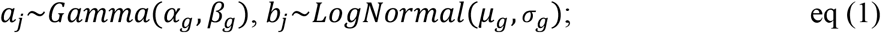

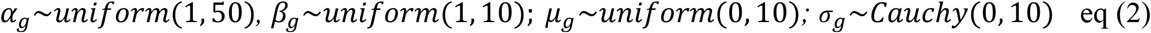

Where *a*_*j*_ and *b*_*j*_ are the parameters of the linear model (parameter *a*_*j*_ complies with a Gamma distribution and parameter *b*_*j*_ complies with a lognormal distribution), *α*_*g*_ and *β*_*g*_ are the parameters of the gamma distribution for parameter *a*_*j*_ of the linear model, *μ*_*g*_ and *σ*_*g*_ are the parameters of the LogNormal distribution for parameter *b*_*j*_ of the linear model. Accordingly, the hyper-parameters *α*_*g*_, *β*_*g*_, and *μ*_*g*_ comply with a uniform distribution and *σ*_*g*_ complies with a Cauchy distribution. Here, the estimated value of parameters *a*_*j*_ and *b*_*j*_ is the median of the simulated posterior sample from the parameters of the linear model estimated with uninformative priors. In equation (1), density functions of Gamma and LogNormal distributions are shown in equations [S8]-[S9], and parameters *α*_*g*_, *β*_*g*_, *μ*_*g*_, *σ*_*g*_ are estimated by PyStan given the estimated median parameter values *a*_*j*_ and *b*_*j*_ of 276,126 biologically plausible and statistically significant probe sets. With the median value of estimated distributions of *α*_*g*_, *β*_*g*_, *μ*_*g*_, *σ*_*g*_ by PyStan, the informative priors are established. The same procedures are applied to the informative prior’s derivation of the rest model parameters.

#### 3.1.2 Prior distributions

The median value histograms of the parameters in continuous models are plotted in Figures S1-S4. Because some extreme values in the posterior sample of parameters *b* and *c* make the distributional shape hard to identify, the 99% confidence level is used for identifying distribution for parameters *b* and *c*. By assimilating these data, the prior distributions of parameters in the continuous dose-response models are summarized in Table 1. Particularly, parameter *a* complies with a gamma distribution. Parameter *b* is symmetrical in the Exponential 2, Exponential 3, Hill, Power, and Linear models, with values larger than 0 complying with a Lognormal distribution. Histograms of the parameter *b* in exponential 4 and 5 models in Figure S2 have two peaks regardless of the size of samples. One peak is within 0-20 and the other peak is within 40-60. A simple model could not describe the complex distribution, so a Gaussian mixture model is selected in Table 1 to describe the shapes. The parameter values of parameter *c* for probes with a decreasing trend in exponentials 4 and 5 are within 0 to 1, and that for genes with an increasing trend in exponentials 4 and 5 is larger than 1. Hence, for genes with a decreasing trend, a beta distribution is used as the parameter range of beta distribution is within 0-1. To keep the consistency, genes with an increasing trend are taken as the 1/beta distribution. The parameter *c* in the Hill model in Figure S3 has a peak around the median parameter value of the uniform prior at ∼7.5. The reason is that the dose-response data for some probes are not sufficient to inform its value with the uniform prior distribution. So, the posterior distribution generated from the Markov Chain Monte Carlo sampler is also almost identical to the prior uniform distribution (i.e., Uniform [0,15]). Consequently, the parameter histogram centers at the median value of the parameter range set by the prior distribution (i.e., the uninformative prior of parameter *c* in the Hill model is [0, 15] with a peak at ∼7.5). When considering the shape of parameter *c* in the Hill model, the peak at ∼7.5 was ignored and a chi-square distribution is used as it is heavy-tailed which takes a higher probability at the large values than other distributions. The same principle applies to parameter *g* in Figure S4. The parameter *g* in exponential 3, 5, and power models also has a peak at ∼7.5 as the uniform prior of parameter *g* is [0, 15]. The parameter *g* of the Hill model shows an obvious increase and a subsequent decrease which fits a chi-square distribution. The chi-square distribution shape of the parameter *g* in the Hill model is referred to and applied to the parameter *g* in exponential 3, 5, and power models to make the informative priors consistent across models.

**Table 1.**
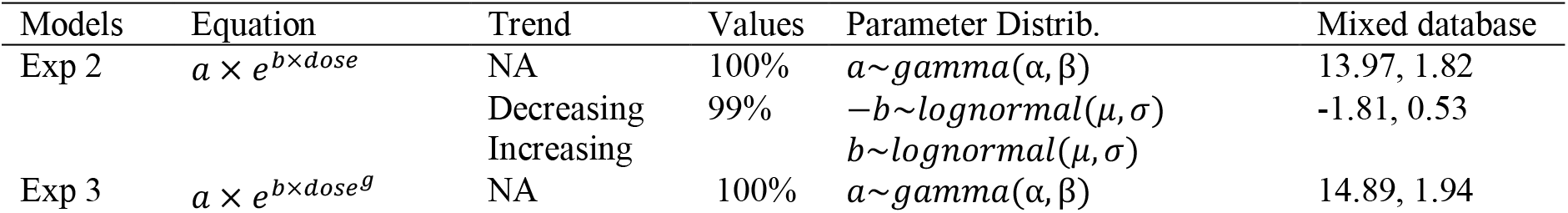
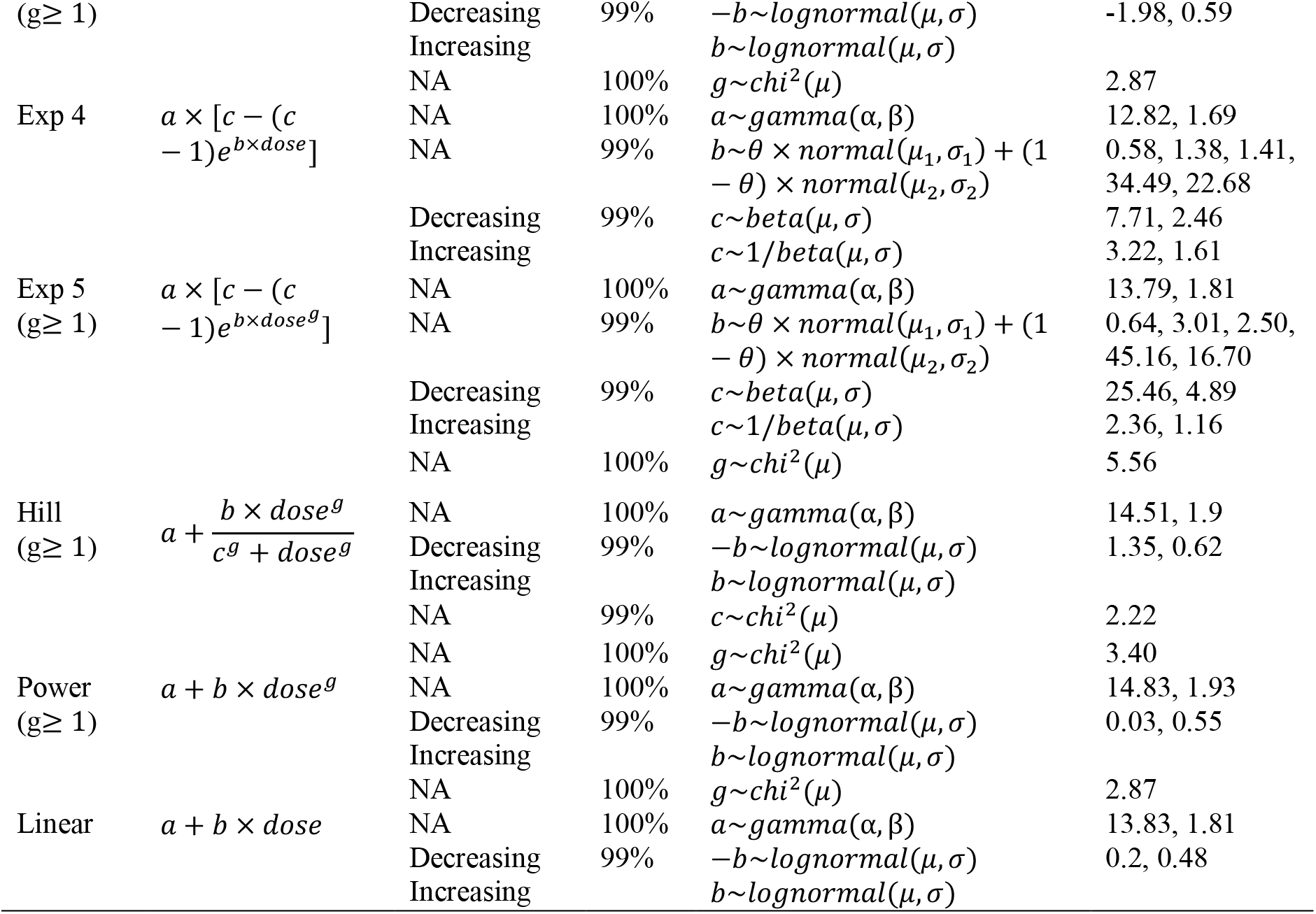
The distribution type and estimated parameters of the informative priors

### 3.2 The performance of informative priors

#### 3.2.1 Effects of informative priors on a single estimate

##### Methods: simulations covering all possible shapes

The identified informative priors in Section 3.1.2 are incorporated with BBMD by PyStan to investigate the performance of informative priors by simulation studies of six “true” genomic dose-response shapes, which are plotted in Figure 2. These six shapes are selected from 100 randomly sampled genes in Figure S5 to reveal the representative dose-response shapes of genomic data. Typical shapes include ‘S’, and ‘J’. Because of its flexibility in reflecting different shapes of dose-response curves, the Hill model is selected to represent the six “true” dose-response shapes with different values of the parameters. The parameter values for the “true” models as well as the parameter of within dose group variance were estimated from real data to represent typical dose-response relationships observed in real genomic data. 1000 samples are randomly drawn from the specified Hill models shown in equations (3)-(5) at 5 fixed normalized dose levels [0, 0.125, 0.25, 0.5, 1.0] with three repeats for each level, where 0 represents the control group and 1.0 represents the maximum dosage. The dose levels are designed based on the typical levels in a toxicological dose-response experiment. The [1000 × 3] vector of **Response** in equation (4) and the **Dose** in equation (5) are input to BBMD to calculate the BMDs under (1) uninformative (i.e., uniform) priors and (2) informative priors derived in section 3.1.2. The real BMD values are derived from equation (6) with BMR (Benchmark Response) defined as one standard deviation shift from the control. The “true” BMD values calculated by equation (6) in the order of chemicals [NPX, cyclosporine, rosiglitazone, TTCP, DNZ, FT] are 44.09, 89.45, 51.91, 60.51, 30.98, 92.20 mg/*kg* ∙ *d* respectively.

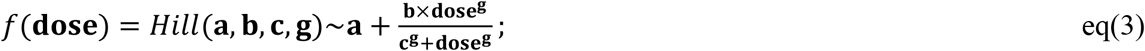

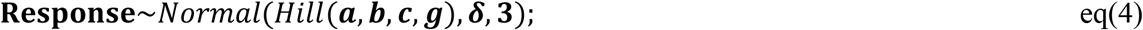

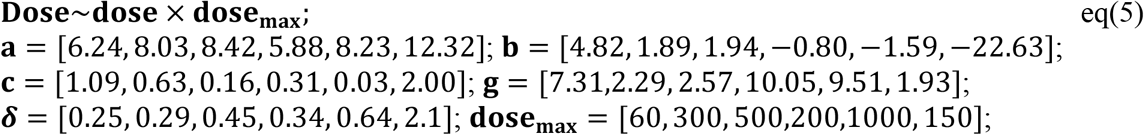

where **a, b, c, g** are the parameters of the specified Hill model, ***δ*** is the designated standard deviation, and **dose**_**max**_ is the maximum dose level of the 6 ‘true’ genomic dose-response data in the order of [NPX, cyclosporine, rosiglitazone, TTCP, DNZ, FT]; dose is a vector of the normalized dose level [0, 0.125, 0.25, 0.5, 1.0]; and **Response** is a vector of 1000 × 3.

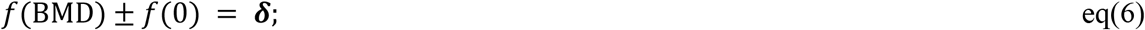

where *f*(0) is the estimated response at zero dose, *f*(**BMD**) is the response at **BMD** for six chemicals, and ***δ*** is a vector of the estimated standard deviation.

**Figure 2.**
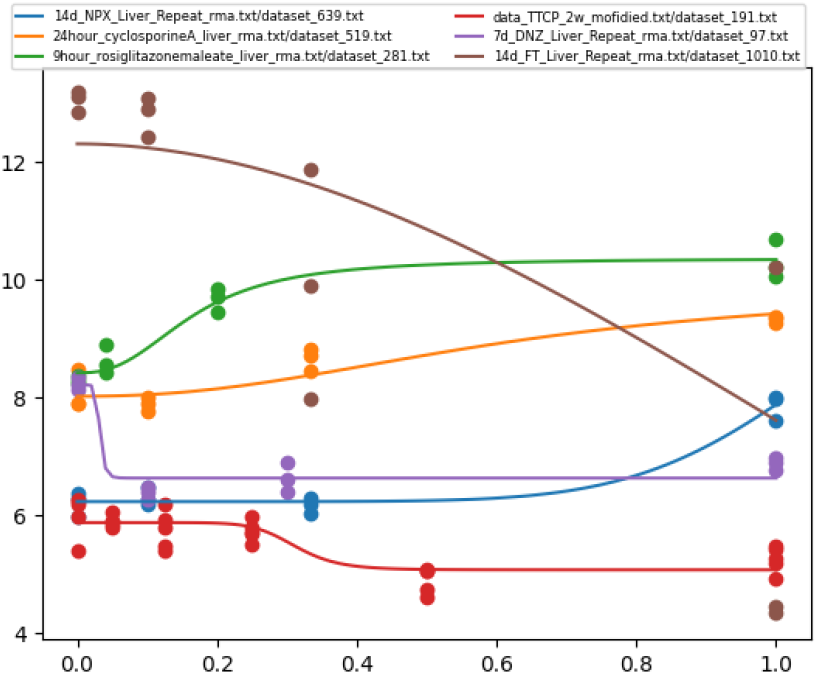
Typical genomic dose-response distributions

The BMDs[uninformative] calculated with uninformative priors and the BMDs[informative] calculated with informative priors are compared with the BMDs[true]. The statistics for single estimate comparison include accuracy rate, the width of 90^th^ percentile interval (reflecting the uncertainty in estimates), and bias comparisons. The accuracy rate is defined as the percentage of BMDs[true] within the 95% confidence interval of estimated BMDs [uninformative or informative] (BMDL-BMDU). For the uncertainty comparison, the ratios of BMDU [the 95% of the BMD] over BMDL [the 5% of the BMD], and the ratios of BMD over BMDL are compared across 7 models and the model-averaged estimates. As a note, BBMD provides the Bayesian model averaging method to combine the estimates of all models based on model weight^10^, i.e., models with better goodness of fit have a higher model weight. In addition, the BMD (or BMDL] estimates are compared with the BMD [true] to reveal the bias difference when using informative and uninformative priors.

##### Results: accuracy rate

The accuracy rates using uninformative and informative priors are summarized in Table 2. When comparing all the 1000 estimates without examining the quality of estimates, only datasets ‘TTCP’ and ‘cyclosporine’ using the informative priors have a slightly higher accuracy rate than the counterparts using uniform priors, while the other four chemicals have a slightly lower accuracy rate when using the informative priors. Typically, the BMD estimates larger than the maximum dose (i.e., the dose-response curve is very flat) are not eligible for dose-response assessment. Simulations with BMD values smaller than the maximum dose level among the original 1000 simulations for the six chemicals are regarded as the valid simulations and their numbers are expressed as N* in Table 2. The corresponding accuracy rates of valid N* simulations (Accuracy*) are also summarized in Table 2. Specifically, datasets ‘TTCP’ and ‘cyclosporine’ using informative priors still have a higher accuracy rate. Besides, the ‘FT’ using the informative prior has a higher accuracy rate but less valid estimates as the N* number of ‘FT’ decreases. BMD estimates with BMD/BMDL > 20 and BMDU/BMDL > 40 indicate that large uncertainty exists. After removing these estimates with large uncertainty from the N* simulations, the remaining simulations are regarded as valid estimates. The number of remaining simulations is expressed as N** with their corresponding accuracy rates (Accuracy**) are summarized in Table 2. The accuracy for the simulations using informative priors increases other than the datasets ‘NPX’ and ‘DNZ’. Except in the case of ‘FT’, BBMD with informative priors provides more valid and less uncertain estimates as the N* and N** values using the informative priors are larger than those using uniform distributions. ‘FT’ is an unusual dataset that the valid simulations represented by numbers N* and N** decrease significantly compared to the original number N after removing unreliable estimates and estimates with large uncertainty. The exception of ‘FT’ may be caused by the inconsistency between the shape of the ‘FT’ dose-response curve and the general dose-response range represented by the informative priors.

**Table 2.**
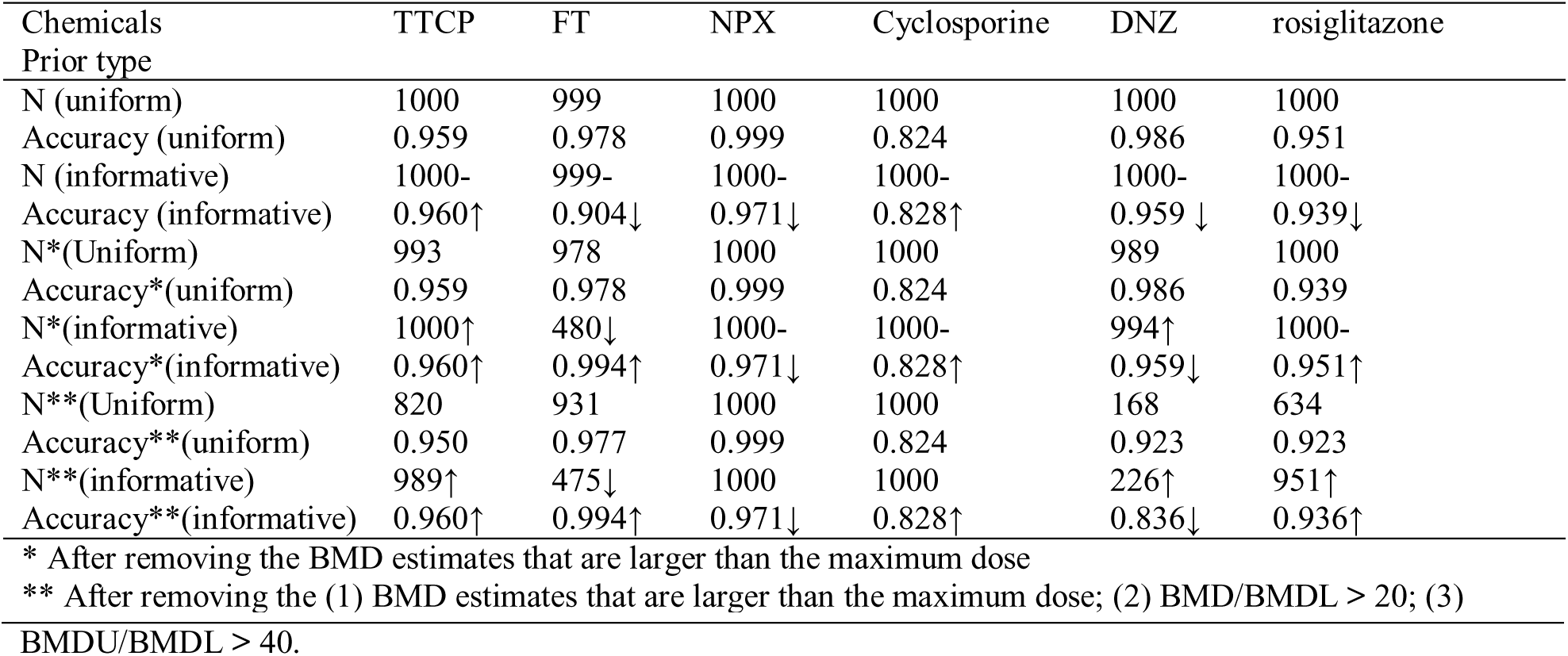
Accuracy rate comparison with uninformative and informative priors

| Chemicals<br>Prior type | TTCP | FT | NPX | Cyclosporine | DNZ | rosiglitazone |
| --- | --- | --- | --- | --- | --- | --- |
| N (uniform) | 1000 | 999 | 1000 | 1000 | 1000 | 1000 |
| Accuracy (uniform) | 0.959 | 0.978 | 0.999 | 0.824 | 0.986 | 0.951 |
| N (informative) | 1000- | 999- | 1000- | 1000- | 1000- | 1000- |
| Accuracy (informative) | 0.960↑ | 0.904↓ | 0.971↓ | 0.828↑ | 0.959 ↓ | 0.939↓ |
| N*(Uniform) | 993 | 978 | 1000 | 1000 | 989 | 1000 |
| Accuracy*(uniform) | 0.959 | 0.978 | 0.999 | 0.824 | 0.986 | 0.939 |
| N*(informative) | 1000↑ | 480↓ | 1000- | 1000- | 994↑ | 1000- |
| Accuracy*(informative) | 0.960↑ | 0.994↑ | 0.971↓ | 0.828↑ | 0.959↓ | 0.951↑ |
| N**(Uniform) | 820 | 931 | 1000 | 1000 | 168 | 634 |
| Accuracy**(uniform) | 0.950 | 0.977 | 0.999 | 0.824 | 0.923 | 0.923 |
| N**(informative) | 989↑ | 475↓ | 1000 | 1000 | 226↑ | 951↑ |
| Accuracy**(informative) | 0.960↑ | 0.994↑ | 0.971↓ | 0.828↑ | 0.836↓ | 0.936↑ |

\* After removing the BMD estimates that are larger than the maximum dose
\*\* After removing the (1) BMD estimates that are larger than the maximum dose; (2) BMD/BMDL > 20; (3)

For the ‘NPX’ with informative priors, there are more inputs that the BMDU values are less than the BMD [true]. ‘NPX’ represents a type of dose-response relationship i.e., response hardly changes until the highest dose level. In this case, the BMD estimates values with informative priors tend to be more conservation than the ones with uninformative priors. In Figure S6, there is no dose-response line that has a sharp change at the 0.8-1.0 dose level. The same phenomenon is observed for the ‘DNZ’, which represents a type of dose-response relationship where response dramatically changes in the low dose region but remains stable afterward. With informative priors, there are more estimates with BMDL larger than the BMD [true], causing the accuracy rate to decrease. Additionally, more BMDL estimates derived from informative prior in the case of ‘FT’ are larger than the BMD [true] when compared with uninformative prior. More than 50% of BMD estimates are larger than the BMD [true] which makes N*=480 after removing estimates of BMD [informative] larger than the maximum dose. The ‘FT’ scenario represents a dose-response relationship with the largest response change consistently across the dose levels. Besides the substantial change in response, the within dose group variance is also very large (i.e., the standard deviation *δ* = 2.1), therefore large parameter ‘b’ value is needed to adequately describe the shape of ‘FT’ in Figure 1. Because it is more likely to sample a large value for parameter ‘b’ from the uninformative prior (i.e., uniform distribution, as long as it is within the bound) than the informative prior, the fitted dose-response curve with an uninformative prior is more flexible to the response with significant change resulting more accurate BMD estimates. In contrast, the rest datasets ‘TTCP’, ‘Cyclosporine’, and ‘rosiglitazone’ have a higher accuracy rate as their parameter values are consistent with the informative distributions.

##### Results: uncertainty

The dispersion of model-averaged BMD estimates across the six chemicals with informative and uninformative priors are measured by various ratios summarized in Table 3. For all chemicals, the 95% confidence interval of ratios ‘BMDU/BMDL’ and ‘BMD/BMDL’ decreases with informative priors. For example, the 95% confidence interval of ‘BMDU/BMDL’ for ‘TTCP’ decreases from 2.50-194.08 to 2.70-14.07. Most median values of ‘BMDU/BMDL’ and ‘BMD/BMDL’ decrease with informative priors except that the ‘BMDU/BMDL’ and ‘BMD/BMDL’ values in ‘FT’ increase from 5.47 to 7.92 and 2.29 to 2.59, respectively, which is still acceptable. The uncertainty reduction in model-averaged estimates with informative priors is evident. On the other hand, we compared the effects of informative priors on the uncertainty of the seven commonly used dose-response models. The uncertainty comparisons of the seven continuous dose-response models for all six chemicals are summarized in Table S.1. As for the ‘BMDU/BMDL’ and ‘BMD/BMDL’ ratios, except exponential 4 and 5 models for ‘DNZ’ have increased intervals, the estimation uncertainty of all other models for all chemicals decreases. The poor performance of exponential 4 and 5 models of ‘DNZ’ was mitigated by the model averaging method for BMD estimation, i.e., the BMD estimates from all models were taken into account with a weight reflecting how well the models fit the data. Although exceptions exist in individual models, the majority of BMD estimates from individual models and all model averaged BMD estimates with informative priors show a decrease in uncertainty.

**Table 3.**
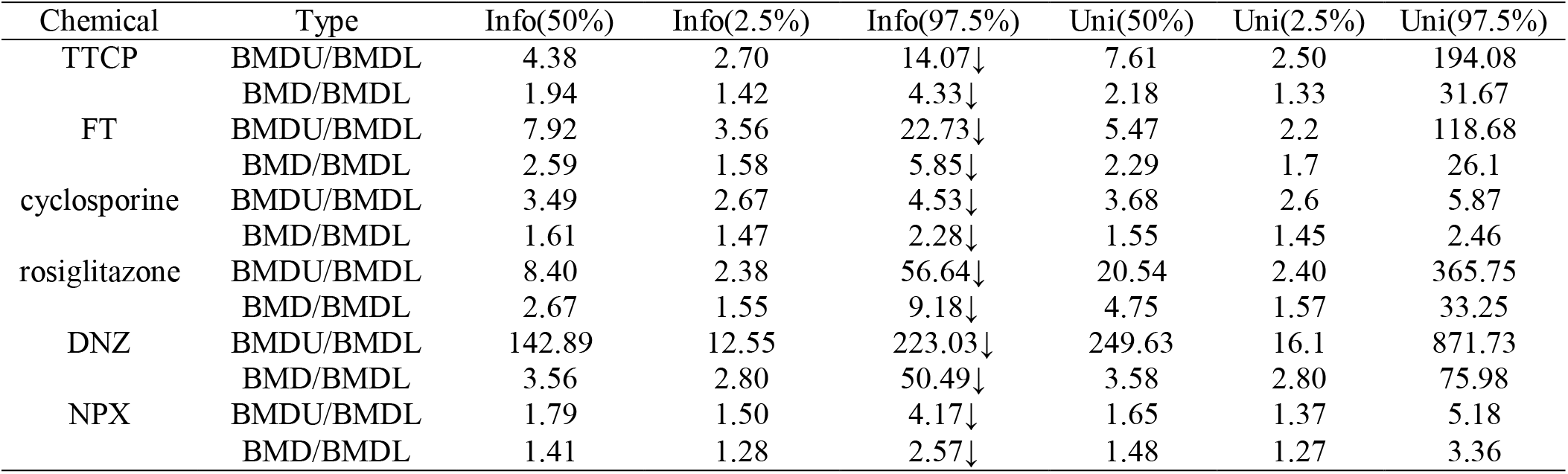
Uncertainty Comparisons of Model Averaging Estimates with Informative and Uninformative Priors

##### Results: bias

The difference between the model-averaged BMD estimates and BMD[true] is summarized as the BMD/BMD[true], and BMD [true]/BMDL ratios in Table 4.When comparing the BMD/BMDL estimates with the BMD[true], first, the majority of the dispersion of the ‘BMD/BMD [true]’ and ‘BMD [true]/BMDL’ ratios decrease. In terms of bias, there is no clear trend how the ratio of BMD/BMD[true] changes for either informative or uninformative priors. For both cases, the median values of BMD/BMD[true] are close to 1 with a range hovers around values from <1 to >1. This indicates that the types of priors have a limited impact on the precision of BMD estimates. The other important indicator BMD[true]/BMDL reveals the conservation of estimations. For both types of priors, the median values of BMD[true]/BMDL are larger than 1 (i.e., most of the estimated BMDL are smaller than the true BMD) indicating that the estimation is conservative and protective. The 2.5% of the BMD[true]/BMDL ratios of ‘TTCP’ and ‘rosiglitazone’ for both types of priors are smaller than 1, and the 2.5% lower bounds for both chemicals increase with informative priors. That is, the BMD estimates for ‘TTCP’ and ‘rosiglitazone’ with informative priors are closer to the BMD [true] and there are fewer BMDL estimates larger than the BMD. While the 2.5% of the BMD [true]/BMDL ratios of ‘cyclosporine’ and ‘NPX’ for both types of priors are larger than 1. That is, for ‘cyclosporine’ and ‘NPX’, both types of priors provide a conservative estimate. In addition, the 2.5% of BMD [true]/BMDL ratios of ‘FT’ and ‘DNZ’ are smaller than 1 with informative priors, while with the uniform priors, these ratios are larger than 1. This explains the decreased accuracy rate of ‘FT’ and ‘DNZ’ with informative priors in Table 2. This also indicates that the informative priors shift the BMD estimates of ‘FT’ and ‘DNZ’ in the right direction and away from the BMD [true].

**Table 4.** Bias Comparisons of Model Averaging Estimates with Informative and Uninformative Priors

| Chemical | Type | Info(50%) | Info(2.5%) | Info(97.5%) | Uni(50%) | Uni(2.5%) | Uni(97.5%) |
| --- | --- | --- | --- | --- | --- | --- | --- |
| TTCP | BMD/BMD[true] | 1.25 | 0.70 | 2.27↓ | 1.27 | 0.69 | 2.97 |
|  | BMD[true]/BMDL | 1.54 | 0.95 | 5.8↓ | 1.73 | 0.93 | 23.39 |
| FT | BMD/BMD[true] | 1.65 | 0.64 | 2.73↑ | 0.91 | 0.37 | 1.60 |
|  | BMD[true]/BMDL | 1.54 | 0.83 | 5.77↓ | 2.65 | 1.21 | 25.74 |
| cyclosporine | BMD/BMD[true] | 0.63 | 0.35- | 1.21↓ | 0.61 | 0.35 | 1.28 |
|  | BMD[true]/BMDL | 2.63 | 1.61 | 4.89↑ | 2.60 | 1.64 | 4.77 |
| rosiglitazon | BMD/BMD[true] | 1.19 | 0.36 | 3.72↓ | 1.15 | 0.23 | 3.88 |
| e | BMD[true]/BMDL | 2.29 | 0.88- | 14.49↓ | 4.86 | 0.88 | 24.19 |
| DNZ | BMD/BMD[true] | 0.85 | 0.35 | 28.45↓ | 0.55 | 0.28 | 29.97 |
|  | BMD[true]/BMDL | 4.18 | 0.45 | 9.22↓ | 6.35 | 1.91 | 12.45 |
| NPX | BMD/BMD[true] | 0.91 | 0.57 | 1.01↓ | 1.09 | 0.58 | 1.15 |
|  | BMD[true]/BMDL | 1.59 | 1.29 | 3.89↓ | 1.36 | 1.13 | 4.27 |

The bias comparison of the seven continuous dose-response models for all six chemicals are summarized in Table S.2. Overall, these bias analysis results agree well with previous accuracy and uncertainty analysis results. The informative priors could reduce the estimate difference between the ‘real’ value and estimates for ‘TTCP’, ‘rosiglitazone’, and ‘cyclosporine’. Although the NPX has a less accuracy rate when using the informative priors in Table 2, the informative priors make the estimates more conservative, which is acceptable on a regulatory basis. But for the ‘FT’ and ‘DNZ’, the informative priors make the estimates less accurate. Therefore, the informative priors may reduce bias when the informative priors are used properly.

#### 3.2.2 Effects on pathway distribution

##### Methods

Based on the National Toxicology Program’s guidance on genomic dose-response modeling^13^, the probes are grouped into significant pathways for BMD estimation. In the BBMD system, the model-averaged BMD estimates previously discussed are further grouped into significant pathways. To investigate the potential effects of informative priors on the pathway BMD distribution, the 24 previously published in-vivo microarray GSE45892^18^ are used. One way ANOVA preprocessing method with the defaults of P-value 0.05 and fold change 2 is used to identify biologically and statistically significant probes. The biological process of gene ontology (GO) in BBMD is used to reveal the pathway difference using the uninformative and informative priors. The cumulative distribution functions (CDFs) of median BMDs are plotted. The Kolmogorov-Smirnov test is applied to compare the distribution differences statistically.

##### Pathway results

The CDF of all biological process GO pathways with the informative and uninformative priors are plotted in Figure 3. All cumulative distributions are similar in shape. The BMD values with informative priors are slightly larger than counterparts using uninformative priors. The P values of the Kolmogorov-Smirnov test for NDPA_5d, MDMB_2w, NDPA_2w, MDMB_4w, NDPA_13w, TTCP_13w, MBMD_13w are all 1.0, which indicates that there is no difference on the seven distributions. The rest distributions have a P value smaller than 0.05 with the range from 0.022 to 2.39e-12, which indicates that the two distributions are statistically different.

**Figure 3.**
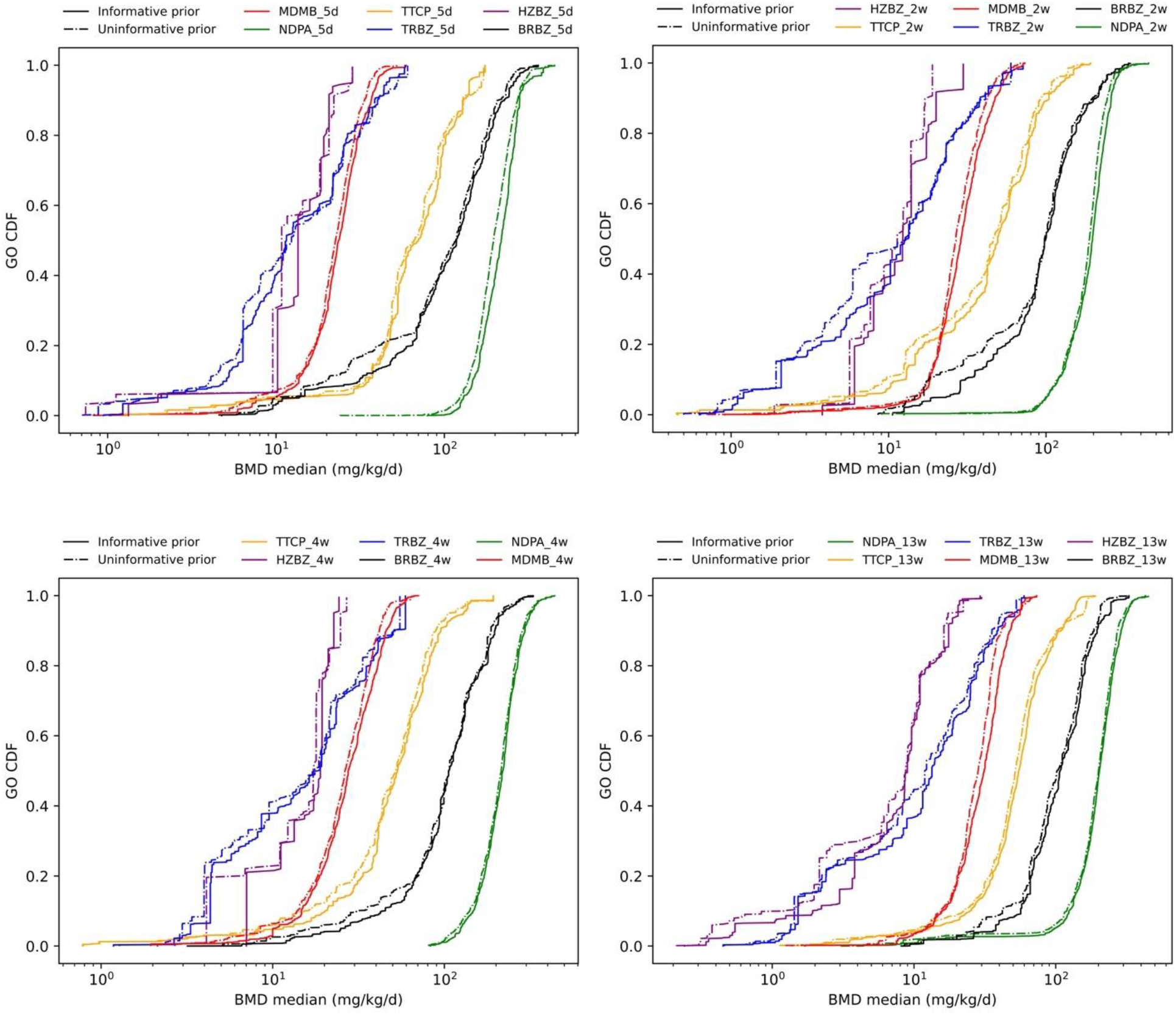
Effects of informative priors on the cumulative distribution function of pathways

#### 3.2.3 Effects on the pathway-based point of departure

##### Methods

Following the pipeline of BBMD, the pathway BMD is eventually used to derive a point of departure value, BEPOD, that could be used for toxicological comparison. Therefore, the 75 molecules data on day 28 in Open TG-Gates archived from^11^ and the previous mentioned GSE45892^18^ are used to compare the effects of informative priors on the correlation relationship between the pathway-based point of departure and the same day apical point of departure. As chemical HZBZ has no apical point for the short term, the HZBZ datasets in GSE45892 are removed for this pathway analysis. In total, 95 datasets are used in this analysis. The same-day apical data as the microarray datasets are gathered from previous papers ^11, 18^. For each chemical, the biological GO pathway results with Fisher’s exact P value ≤ 0.05 and percentage > 2% are kept as the enriched pathway. The median BMD(BMDL) value of the enriched pathway is used as the biological pathway-based point of departure, BEPOD. The correction relationship between the pathway-based BMD (BMDL) and the apical point of departure BMD (BMDL) are compared respectively under two situations: (1) with uninformative priors (i.e., uniform distribution) and (2) with informative priors. Criteria including Pearson’s correlation coefficient and the root-mean-square deviation (RMSD) are used. To capture the relative concordance of the pathway-based point of departure and the apical point of departure, the two values are transformed to the log scale with base 10^11^.

##### Pathway-based POD results

The correlations between the pathway-based point of departure and apical point of departure are plotted in Figure 4. Based on our previous case studies^10^, we choose the median pathway BMD (BMDL) values to represent the pathway-based point of departure values as the median pathway BMD (BMDL) values have a high correlation with the apical point of departure BMD (BMDL) values. An empirical ratio factor, 0.24-0.25, is found by linear regression between the pathway-based BMD (BMDL) and the apical BMD (BMDL). An empirical ratio of 0.11 is applied to the pathway-based BMD and the apical BMDL. It is important to note that the empirical ratio does not affect the correlation relationship. The ratio factors are applied to scatterplots with both informative and uninformative priors plotted in Figure 4. The Pearson’s correlation coefficients with informative priors range from 0.77-0.80, and the *r* with uninformative prior is 0.72-0.79. Overall, the pathway-based point of departure is highly correlated with the apical point of departure. The RMSD is 0.49-0.55 with informative prior and is 0.50-0.66 with uninformative priors. Results indicate that the correlation slightly increases and the RMSD slightly decreases with the informative prior.

**Figure 4.**
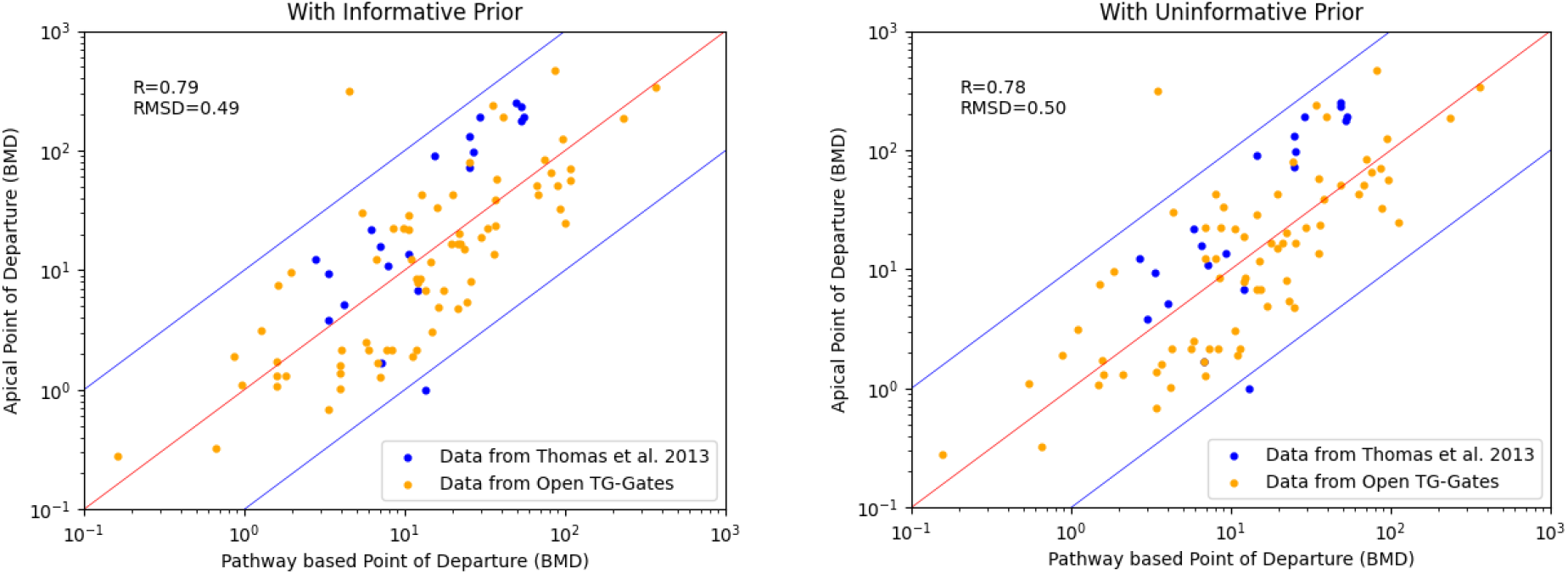
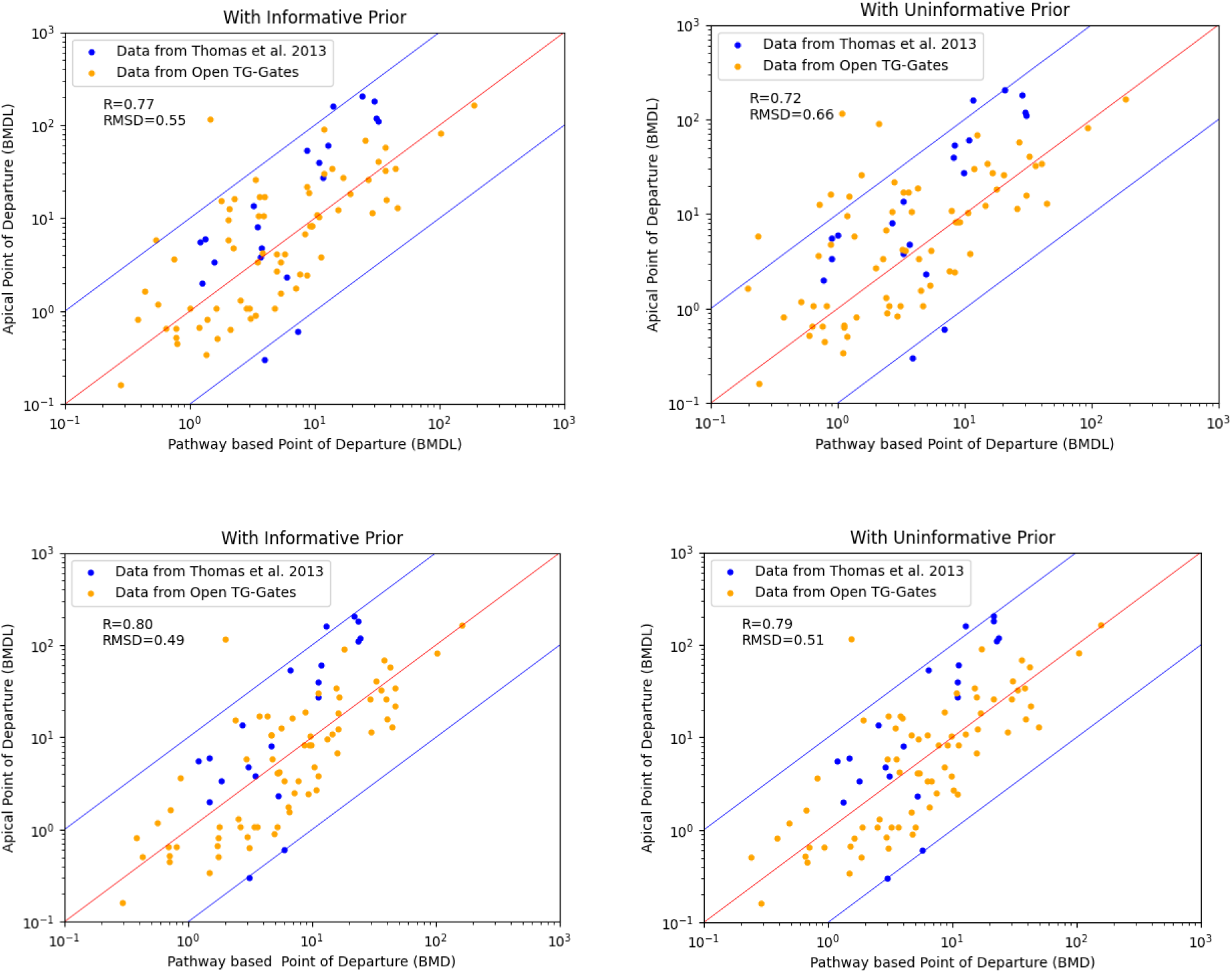
Effects of informative priors on the pathway-based point of departure

## 4. Discussion

This paper describes the process of deriving informative priors for the commonly used continuous dose-response models for genomic data. The distributions of parameters *a* in all continuous models and *b* in the Exponential 2, 3, Linear, Hill, and Power models are quite certain. But the parameter *b* in the exponential 4 and 5 models in Figure S2 is hard to describe using a single distribution as there are two peaks, which are permanent as their shapes do not change regardless of the size of the data. The other peak for both Exponential 4 and 5 models concentrates at the value of 50. The peak at the value of 50 could not be ignored as it represents the parameters for probes with a large response change. A compromised solution applied in this manuscript is to use a Gaussian mixture model to allow sampling from both peaks, but this approach still has the risk to sample inadequate value from the prior distribution. So, a more elegant method would be to have a gene level informative priors in which the values of their parameters are derived from the individual probe data. Another obstruct to deriving the informative priors is the noise of parameter estimates. As most of the genomic dose-response data used in this study for deriving the informative priors only have four dose levels and three repeats (data from Open TG-Gates), it is challenging to get robust parameter estimates for models with four parameters. Without sufficient information, the posterior parameters are centered around the median value of the prior uniform distribution, which provides almost no information on what the real parameter value is (e.g., the parameter *c* in the Hill model and parameter *g* in Exponential 3, 5, and Power models). After removing the probe sets with limited information for characterizing parameter value, a heavy-tailed distribution is more appropriate to describe the distributional shape because larger values are associated with higher probability than other distributions. This simplified method may not reveal the real distribution, more informative genomic datasets may be helpful to overcome this limitation. Additionally, deriving informative priors at the individual gene level may be another possible solution.

The effects of informative priors on the individual genomic dose-response level and the pathway level were investigated. In terms of individual levels, the informative priors work well with the ordinary/common distributions and could improve the accuracy, decrease the bias, and reduce the uncertainty. But when dose-response relationship is not consistent with the shape represented by the informative prior (e.g., a dramatic change in response only occurs in very high or low dose levels), informative priors can have significant impact on fitted dose-response curve. (i.e., reflecting common situations) which may lead to to a decrease in accuracy. Alternatively, instead of using a generalized and fixed informative prior, an individual gene based informative priors may be a better choice, i.e., informative priors are derived for each probe. That is, the parameters of informative priors change along with the dose-response shape of individual genomic dose-response data. This will be an exceedingly computationally intensive approach and require further research. In addition, the informative priors have limited effects on the pathway distribution as each pathway has a group of genes. The estimates using either the informative priors or uninformative priors are quite stable on a large scale with thousands of genes and both results are acceptable. The CDFs of the BMD median pathways are similar but with a slightly increased BMD estimate. The correlation between the pathway-based point of departure and the apical point of departure on the same time scale increases slightly with the informative priors than with the uninformative priors, especially for the correlation of BMDLs. A possible reason is that the reduced uncertainty resulting from informative prior makes the BMDLs closer to BMDs which have a higher correlation coefficient.

Generally, the informative priors can reduce estimation uncertainty and bias, and improve accuracy and correlation. At the individual gene level, generalized informative priors may not be adequate in every case, so extra attention is required when generalized informative priors are applied. Further research on more elaborate informative priors is needed. At the pathway level, informative priors are quite stable as the uninformative priors but provide a slightly improved correlation between the pathway-based point of departure and the apical point of departure.

## 5. Conclusion

Our work empirically derived the informative priors based on current available genomic dose-response data, and systematically tested the effects of informative priors on the genomic BMD estimates at both the individual level and pathway level. When appropriately applied, informative priors will benefit the estimates by improving accuracy and reducing uncertainty. This work provides powerful incentives to use empirical data-derived informative priors in dose-response modeling for continuous genomic data to derive a pathway-based point of departure which is better correlated with the counterpart estimated from apical endpoints.

## Acknowledgments

Research reported in this publication was supported by the National Institute of Environmental Health Sciences of the National Institutes of Health (NIH) under Award Number R42ES032642. The content is solely the responsibility of the authors and does not necessarily represent the official views of the National Institutes of Health.

